# Tumor spheroids accelerate persistently invading cancer cells

**DOI:** 10.1101/2022.04.04.486939

**Authors:** Melanie Audoin, Maria Tangen Søgaard, Liselotte Jauffred

## Abstract

Glioblastoma brain tumors form in brains’ white matter and remains one of the most lethal cancers despite intensive therapy and surgery. The complex morphology of these tumors includes infiltrative growth and gain of cell motility. Therefore, various brain-mimetic model systems have been developed to investigate invasion dynamics. Despite this, exactly how gradients of cell density, chemical signals and metabolites influence individual cells’ migratory behavior remains elusive. Here we show that the gradient field – induced by the spheroid – accelerates cells’ invasion of the extracellular matrix. We show that cells are pushed away from the spheroid along a radial gradient, as predicted by a biased persistent random walk (BPRW). Thus, our results grasp in a simple model the complex behavior of metastasizing cells. We anticipate that this well-defined and quantitative assay could be instrumental in the development of new anti-cancer strategies.

## Introduction

Glioblastoma cancer is the most common type of primary, malignant brain tumor in adults. Its high mortality rate is accredited to its aggressive invasion of the surrounding healthy tissue. Prior to invasion, epithelial (tissue-like) cancer cells gain motility and leave the primary tumor to invade and ultimately form secondary tumors elsewhere in the organ (or the organism). This transformation from epithelial to mesenchymal (fibroblast-like) is necessary for glioblastoma cells to gain motility. Mesenchymal motility – or *crawling* – is associated with elongated cell shapes, reinforcement of the intracellular actin network and strong interaction or modification of the local microenvironment^1, 2^. Moreover, the mesenchymal phenotype has been linked to augmented therapy resistance^3^. Although glioblastoma cells are known to migrate via mesenchymal migration modes, amoeboid (leukocyte-like) migration has also been recorded; perhaps as a way to further confer therapy resistance^4^. Hence, cancer cells can move in different modes – either individually or in a variety of configurations – and can switch between them in response to their environment^5, 6^.

The motility of cancer cells has been studied in various *in vitro* tumor models grown from cancer cell lines (or tumor tissues) in extracellular matrix (ECM) or hydrogel environments, which enable the investigation of spheroid invasion and migration in a natural, yet controlled manner^7, 8^.

Previous studies have shown that the most effective direction of motility is outwards and away from the spheroid^9^. Hence, we wondered whether such migrating behavior is intrinsic to the individual cells or whether it is imposed by some external gradient in the vicinity of the tumor spheroid? As the motility of cancer cells in ECM can be described as a persistent random walk (PRW), we hypothesized that the spherically symmetric gradient – imposed by the geometry of the spheroid – gives rise to a bias which drives cell invasion along the radial vector. That is, the cell trajectories would follow a biased PRW (BPRW). To investigate this hypothesis, we tracked glioblastoma (U87-MG) cancer cells embedded in 3D ECM migrating either i) individually or ii) from multicellular spheroids. These experiments were complemented by simulation of trajectories in both assays. Our results confirmed that indeed spheroids exert a repulsive gradient force which pushes the cells to migrate faster, straighter, and radially away from their origin, as predicted by a BPRW.

## Results

Through precise cell tracking of human glioblastoma cells (U87-MG), we characterized the motility patterns of cells in 3D ECM consisting of 65% Matrigel™ in cell culture medium. With the aim of comparing the metastatic spread of cells from tumor spheroids to the migration of individual cells, we prepared two assays as sketched in fig. 1A.

**Figure 1.**
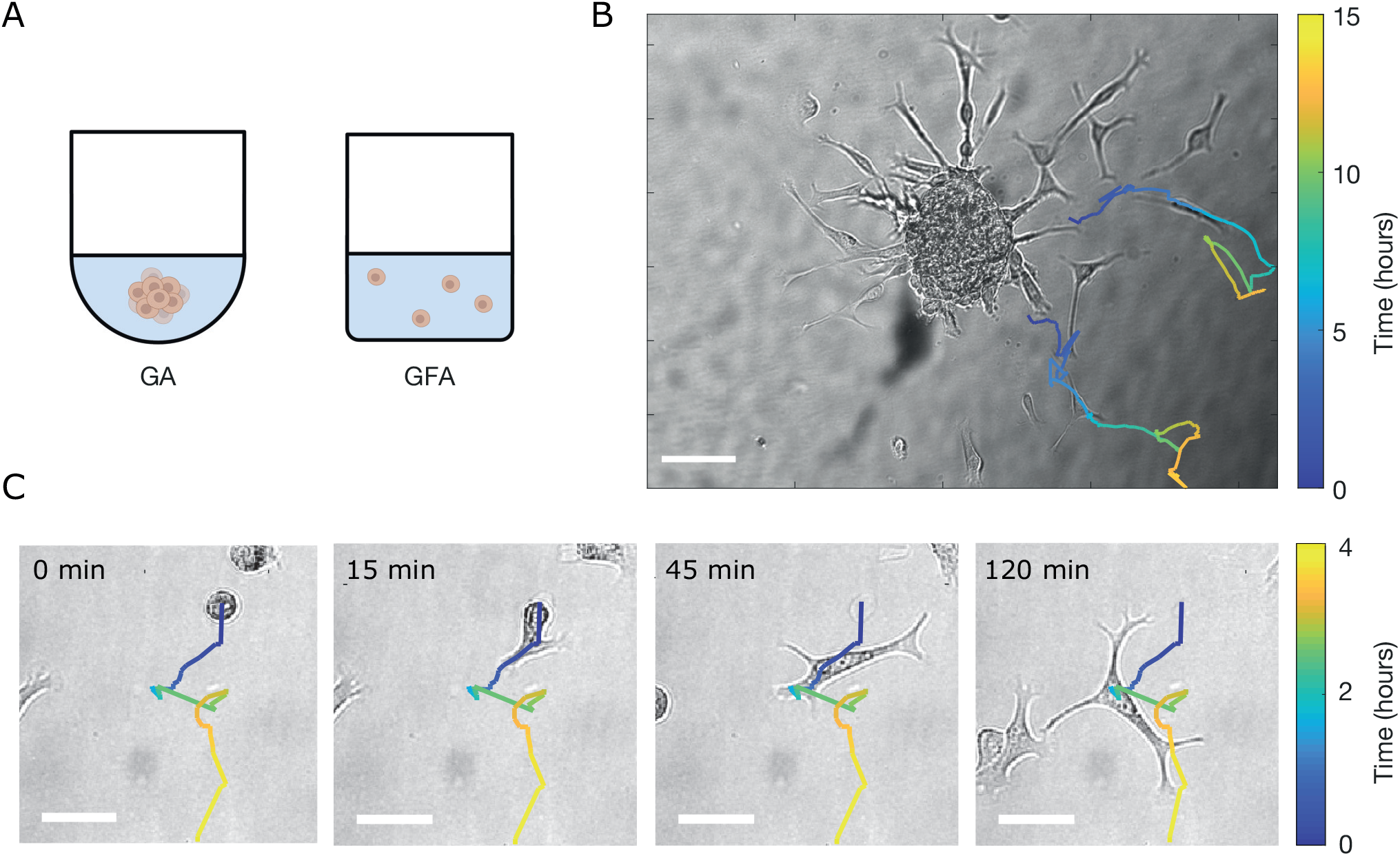
Brain cancer cells invading the ECM. **(A)** Schematics of an ECM-embedded spheroid in a gradient assay (GA) or individual cells in a gradient-free assay (GFA). **(B)** Tracks of migrating cells in a GA from *t* = 9hours and *t* = 5hours, respectively, superimposed on the corresponding bright field image. For each track, the color bar indicates time evolution. Scale bar is 100 μm. **(C)** Morphological transition of a cell prior to migration. Image sequence of brain cancer cells invading the ECM at *t* = {0,15,45,120} min in GFA. The track of the cell is superimposed on the bright field image. The color bar indicates time evolution and the scale bars are 50 μm.

First, spheroids were formed as a result of gravity-assisted accumulation of cancer cells at the bottom of U-bottom wells. After harvesting, the spheroids were embedded in ECM and incubated under physiologically relevant conditions. Then after a few hours, multicellular strands invaded the surrounding matrix and cells gradually left these strands to migrate individually. Representative snapshots of cells superimposed with trajectories are shown in fig. 1B, where cells appear to move radially outward. Due to the spheroid-induced gradient, we refer to this as a gradient assay (GA).

In parallel, a gradient-free assay (GFA) was prepared by embedding non-spheroidal cell cultures in 65% Matrigel™. Fig. 1C shows representative bright-field images of an epithelial-to-mesenchymal transition of a single cell overlaid with the full cell trajectory. The cells were still round-shaped and immobile when first embedded in Matrigel™ at *t* = 0 min. However, with time (1-3 hours) they switched to more elongated shapes with long lamellipodia - characteristic of mesenchymal motility^4^- and started migration.

### Spheroids induce stronger super-diffusivity

A classical metric for characterization of diffusing entities is the anomalous exponent, *α*, which can be obtained from fitting the ensemble-averaged mean-squared displacements, msd (eq. 10), with a power law:

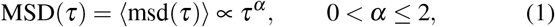

where *τ* is the time lag and 〈…〉 signifies an ensemble average. *α* = 1 corresponds to freely diffusing cells i.e. Brownian motion and in 2D MSD(*τ*) = 4D*τ*, where *D* is the diffusion coefficient with dimensions of m^2^/s. Impeded or constricted motion – as one would expect for non-motile cells in a dense material such as ECM – is sub-diffusive with *α* < 1, whereas super-diffusive motion or active cellular transport is characterized by 1 < α ≤ 2. α = 2 corresponds to ballistic motion.

MSD curves for GA and GFA are shown in fig. 2A where the inset shows the average *α*’s with 95% confidence interval for the two different assays. The anomalous coefficients are *α* = 1.397 [1.384 - 1.410] and *α* = 1.315 [1.299 - 1.330] for GA and GFA, respectively. Hence, in both cases cells are super-diffusing in accordance with prior findings^10, 11^, but disjoint confidence intervals prove that the two experiments are significantly different. Thus, we conclude that cells migrating from a spheroid diffuse faster than cells migrating individually. This result is indicative of a repulsive acceleration of cells in the spheroid periphery.

**Figure 2.**
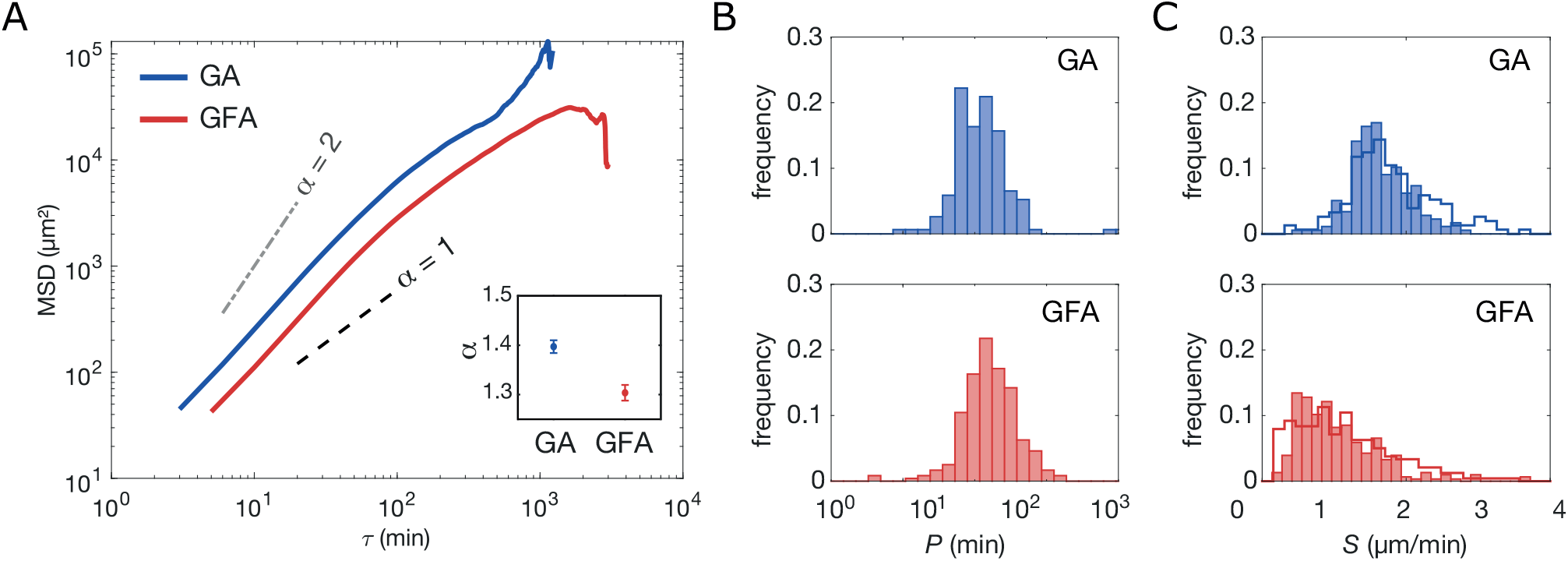
Ensemble-averaged mean-squared displacement analysis. **(A)** GA (N=177) and GFA (N=305) MSD(*τ*) (mean ± SEM) on double-logarithmic scale. Dashed lines corresponding to diffusive (*α* = 1) and ballistic motion (*α* = 2) are inserted as a guide to the eye. Inset: The fitted anomalous coefficient, α, with 95% confidence intervals. **(B)** Distributions of persistence times, *P* on a log-normal scale, of the cells in GA (*N* = 156 with *R*^2^ > 0.99) and in GFA (*N* = 251 with *R*^2^ > 0.99). Values are obtained from the msd-analysis described in eq. 2. **(C)** Distributions of migration speed, *S*, for GA and GFA, both fitted from Fürth’s formula (full line) as in (B) and from averaging over the individual trajectories (bars) calculated from eq. 3.

At long delays (*τ* > 200min) the ensemble-averaged MSDs no longer obey the power law (eq. 1) and the cells lose their super-diffusivity over time. These variations might be due to cell-cell adhesion or pauses during mitosis. Additionally, the mesh size of the extracellular matrix is not perfectly homogeneous which is likely to influence cell motility and accordingly the persistence of the motion. Moreover, the MSD values for *τ* > 800 min suffer from large uncertainties due to the sparse number of data points. Hence, we cannot conclude on these apparent drastic changes at long delays.

### Cells migrate with persistence

Metastatic cancer cells – like many other motile cells – actively move with persistence. This behavior is often modeled by a random walk (RW) with memory, termed a persistent random walk (PRW). A cell moving with persistence is more likely to keep moving in the same direction than in any other. Therefore, any given trajectory can be identified by a characteristic time termed the persistence time, *P,* using Fürth’s formula for a 3D cell trajectory imaged in 2D:

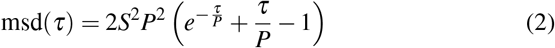

where *S* is the average speed over the entire trajectory and *τ* is still the time delay.

We fitted all GA (blue) and GFA (red) cell trajectories’ msd (eq. 10) by Fürth’s formula (eq. 2) and found the distributions of *P* and *S* shown in fig. 2B and 2C, respectively. The *P*-distributions are log-normal and similar, with most *P* < 90 min: For GA 〈*P*〉 = (12.7 ±1.0) min (mean+SD) of logarithmic values, and for GFA 〈*P*〉 = (16.6 ±1.0) min of logarithmic values. However, we do observe a few highly persistent cells in GA as earlier reported^9^. As these highly persistent cells are only found escaping spheroids (GA), we suggest that this is more likely to be related to local chemical gradients or remodeling of the extracellular matrix rather than genotype or other cell-intrinsic factors. The PRW model assumes that cells migrate with a constant persistence and we thus estimate an average *P* for the entire trajectory even if fig. 2 indicates that cell motility modes change at long time-scales^12^.

The effect of the spheroid on cell motion can be further characterized by the distributions of cell speed, *S*, shown in fig. 2C, which were obtained in two ways: either through msd-fitting with eq. 2 (line) or by averaging over the individual trajectories (bars) as shown below

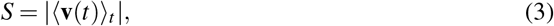

where 〈…〉_*t*_ indicates the average over all *t*’s of a trajectory. The distributions of *S* in fig. 2C exhibit a noticeable right-skew in the GFA compared to GA. Moreover, a Kolmogorov-Smirnov test rejects the null hypothesis that the *S*-distributions are normal distributions.

From fits eq. 2, we found that for cells in GA, 〈*S*〉 =(1.81 ±0.66) μm/min (mean+SD) and in GFA, 〈*S*〉 = (1.15 ±0.71) μm/min (summarized in table 1). Thus, the spheroid seems to induce a repulsion that drives cells persistently and fast away from it.

**Table 1.**
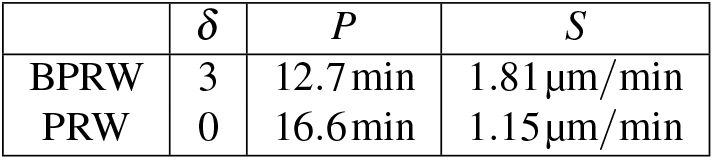
Simulation parameters.

### Cells do occasional U-turns

The angular displacement is defined as the magnitude of the angle, *θ* (*τ*), between two velocity vectors separated by the delay *τ*, i.e., between **v**(*t*) and **v**(*t* + *τ*):

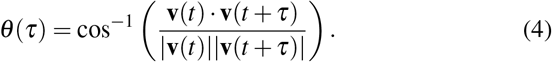

The distributions of *θ*(*τ*) in Supplementary fig. S1 show that cells (*τ* = 15 min) will most often continue in the same or a very similar direction (small *θ*) for both the GA and GFA. This behavior reveals that for short delays, successive velocity vectors are highly correlated. However, for longer *τ* the distributions flatten which indicates that the cell motion gets increasingly uncorrelated with time.

It is worth noticing that for all *τ*’s, *θ* peaks at 0° and 180°. This indicates an anisotropy in the cell migratory path and proves the existence of an axis along which the cells preferably migrate forward and backwards. As mesenchymal cells proteolytically degrade the matrix when moving through it^13–15^, they will most probably use already degraded ECM tunnels when available – either straight ahead or by making a U-turn – as found earlier for human fibrosarcoma in agarose (GFA)^16^ and for U87-MG in a similar assay (GA)^9^.

### Cells leave spheroids by moving fast along a radial gradient

As our goal was to capture the differences between the two assays, we developed a set of descriptive and fairly simple models to complement our experimental data. We used a continuous stochastic model to simulate cell trajectories, where a cell was represented as a single point corresponding to its center-of-mass and its 2D projected trajectory **x**(*t*), which consisted of *n* steps of size *dt*. In particular, we focused on a PRW model as cell migration has been reported to be well described by this^16^. We hypothesized that the gradients imposed by the spheroid would result in an apparent bias in the cell trajectories i.e. following a BPRW model, as previously suggested^17^. Specifically, we anticipated this bias to be a repulsive field in the direction of the spheroid and decaying as 1/ |**r**|, where **r** is the radial vector. The model is described in details in the Materials and Methods section and parameter values are summarized in table 1.

We thus turned to a quantitative description of the differences observed in fig. 1, in particular regarding the direction of motion. To do so, we defined the following two vectors: the major axis of motion, **m**, and the radial vector, **r**. The first is

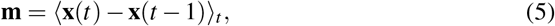

where 〈…〉_*t*_ signifies the average displacement vector of the trajectory, i.e., averaged over all *t*’s. The radial vector is

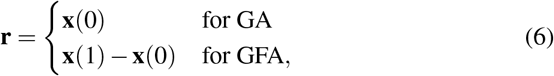

where the position vector **x**(0) equals the radial vector from the center of the spheroid, (0,0), to the initial position on the spheroid surface, as sketched in the inset of fig. 3 (GA). As GFA lacks radial symmetry, we defined the radial vector, **r**, in this assay to be the displacement vector of the first step of the trajectory (see inset of fig. 3). Given this, *φ* is the angle between the vectors **m** and **r**. The *φ*-distributions for all trajectories in GA (blue) and GFA (red) are given in fig. 3, where the dashed lines correspond to the distributions of simulated trajectories. To a great extent, cells in the GA environment follow the radial vector away from the spheroid (*φ* < 75°). In contrast, cells in the GFA do not migrate along any predefined direction which renders the *φ*’s evenly distributed. Furthermore, this is reflected in the medians, which are 35° and 71° for the GA and GFA cells, respectively. Hence, these results suggest that the spheroid repulses cells in the radial direction specifically.

**Figure 3.**
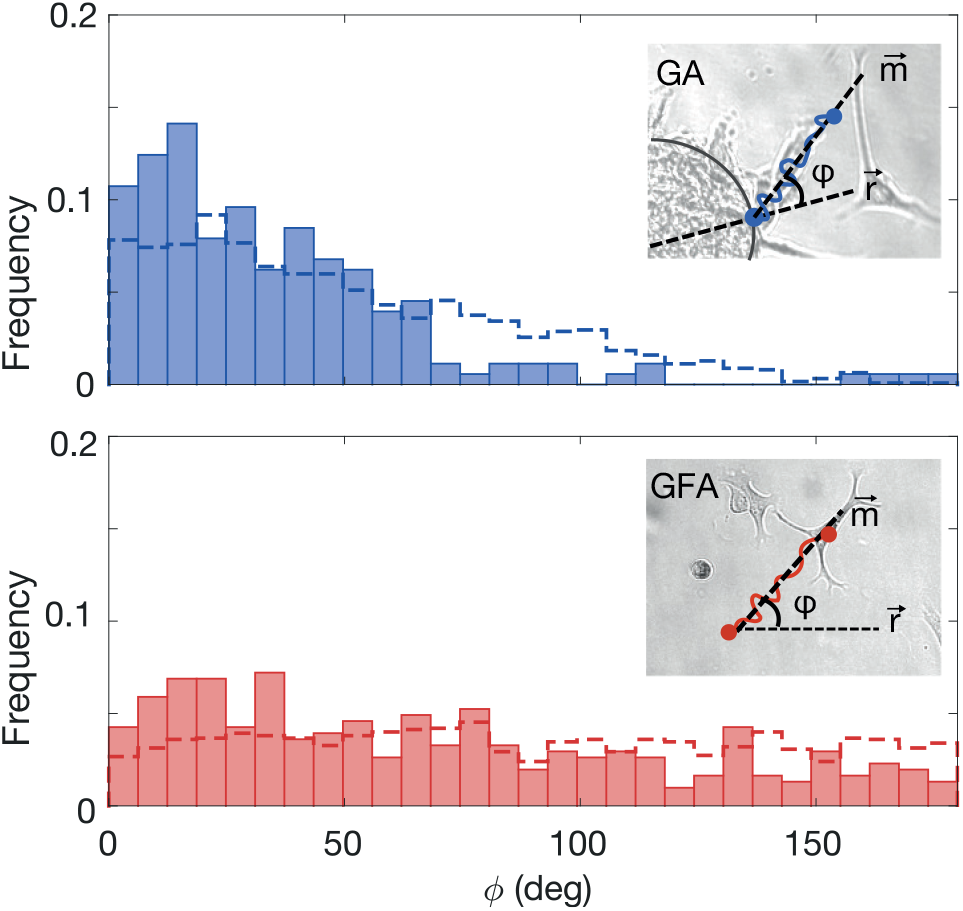
Distribution of angle, *φ*, between the major axis, **m**, and initial radial direction, **r**, for data (bars) and simulation (dashed line) from GA (N = 177) and GFA (N=305). Insets: Schematic of the angle φ between major direction of motion, **m**, and initial radial direction, **r** for GA and GFA, respectively.

### Cells migrate with high directionality away from spheroids

To investigate this further, we calculated the time evolution of the radial distance traveled by the cells:

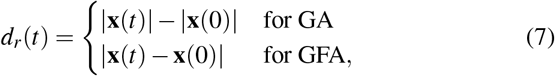

as sketched for both assays in the insets of fig. 4A. The time evolution of the ensemble-averaged radial displacement, 〈*d_r_*(*t*)〉, is also shown in fig. 4A. At any given time point, the GA cells have on average migrated further from their starting point than cells in the GFA. This is in agreement with the observation that GA cells migrate faster than cells in the GFA (fig. 2). It is further justified by the 〈*d_r_*(*t*)〉 obtained from simulated trajectories of both BPRW (blue dashed line) and PRW (red dashed line). To understand this behavior further, we show the radial distance (eq. 7) of individual trajectories in Supplementary fig. S2 and find that he smooth ensemble averages of *d_r_*(*t*) covers very noisy individual trajectories.

**Figure 4.**
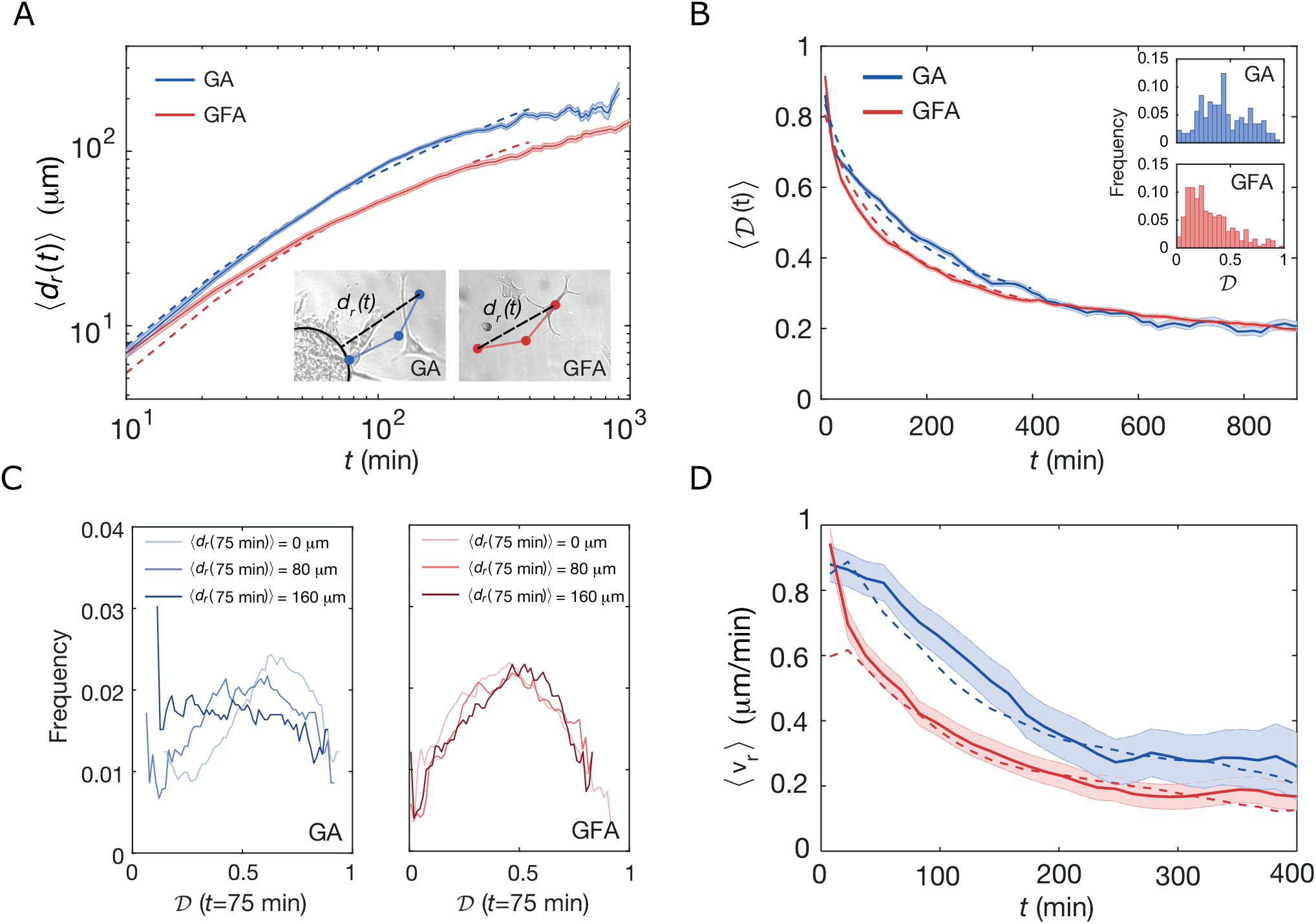
Time evolution of migration patterns. **(A)** Time-dependent ensemble-averaged radial displacement, 〈*v_r_*(*t*)〉, versus time, *t*, (mean ± SEM) for GA (N=177) and GFA (N=305) on a double-logarithmic scale. Dashed lines correspond to results obtained from BPRW (blue) and PRW (red) simulations. Insets: Sketches defining the radial displacement, *d_r_*(*t*), for GA and GFA, respectively. **(B)** Time evolution of the ensemble-averaged directionality, 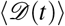 (mean ±SEM) for GA (N=177) and GFA (N=305). The dashed lines are averages over simulated tracks for BPRW (blue) and PRW (red). Insets: Distribution of the directionality of the entire trajectories, 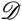. **(C)** Distribution of directionalities of the trajectories after *t* = 75 min, 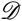(75min), at different distances from initial position, 〈*d_r_*(75min)〉, for GA (blue) and GFA (red). **(D)** Time-evolution of the ensemble-averaged radial velocity, 〈*v_r_*(*t*)〉, (mean ± SEM) for GA and GFA and simulated trajectories (dashed lines) from BPRW (blue) and PRW (red). In all four cases, 〈*v_r_*(*t*)〉 was binned over 15 min with a moving average of 10 data points.

We find that the distance traveled by a cell also depends on the time-dependent directionality, 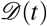. It is defined as the ratio between a cell’s distance from its initial position, **x**(0), and the trajectory length at time *t*:

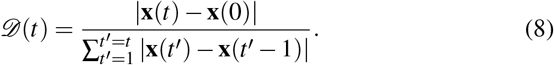

So by definition 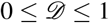, where 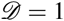 corresponds to a ballistic trajectory and 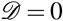 is an infinitely long and tortuous random walk. Fig. 4B shows the ensemble-averaged directionality, 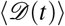, for both GA (blue) and GFA (red) cells as well as for the simulated trajectories (dashed lines). For *t* < 400 min, cells in GA exhibit higher directionality than GFA cells, which proves that cells in GA indeed move straighter. However, at longer time scales, the GA 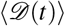 falls off to values closely matching those of cells in GFA. This confirms the idea of a repulsive field around the spheroid with a gradient of −1/**r**^2^. It follows that once the GA cells escape this field, they slow down and their migration patterns start to resemble those of cells moving in a gradient-free environment. The insets in fig. 4B show distributions of directionalities for the entire trajectories, 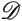. As expected – and despite large variations – cells in GA move with higher overall directionality than cells in GFA.

To test the prediction that the bias field is dependent on the distance from the spheroid center, (0,0), we examined how distributions of 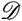 changed at *t* = 75 min with *d_r_*(75min) = {0,80,160} μm as shown in fig. 4C. For GA, the distributions shift towards lower 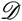 as *d_r_*(*t*) increases, in accordance with the idea that the gradient is strongest close to the spheroid (*d_r_*(75min) =0 *μ*m vs. *d_r_*(75min) = 160μm). In contrast for cells migrating in GFA, there are no spatial changes of *d_r_*(*t*) = {0,80,160} *μ*m as all of them overlap. These results suggest that in the vicinity of the spheroid, cells migrate more efficiently through the ECM. On the other hand, in GFA cell motion is more diffusive so the cells explore more thoroughly their local environment. This means that the cells’ migration strategy is highly environmentally dependent and is balanced between local exploration and a more directed migration mode.

### Spheroids accelerate the radial velocity

We have now established that metastatic glioblastoma spheroid cells invade their surrounding environment faster, more persistently, and straighter than single cells in the exact same ECM. However, a potential time- and space dependence of the proposed bias still needs to be investigated. As the gradient is anti-parallel with the radial vector **r**, we measured the time and space dependence of the radial velocity i.e. the scalar projection of **v**(*t*) onto the unit radial vector:

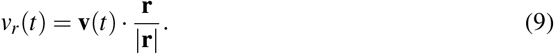

In the GA, the ensemble-averaged radial velocity, 〈*v_r_*(*t*)〉, decays with respect to the radial distance, *d_r_*(*t*), for the first ~ 100 μm after leaving the spheroid (Supplementary fig. S3) after which the spheroid-induced field becomes negligible and the velocity reaches 〈*v_r_*(*t*)〉 ~0.4μm/min. This decay in velocity is, thus, a manifestation of the space dependence of the bias induced by the spheroid. In contrast, in GFA there is no such dependence. Thus, cells in this assay migrate with constant 〈*v_r_*(*t*)〉 ~ 0.2μm/min through a seemingly isotropic space.

Furthermore, 〈*v_r_*(*t*)〉 decays with time for both GA and GFA as shown in fig. 4D. However, for the first part (*t* < 200 min) the radial velocity of GA is significantly larger than in the GFA. The agreement between experiments and simulations confirms that the spheroid-induced field is well described by a – 1/ |**r**^2^| gradient and the bias term of eq. 16.

## Discussion and conclusion

Here, we report migration patterns of glioblastoma U87-MG cells in brain-mimetic culture models. We showed that cells fleeing a spheroid tumor model (GA) move with persistence, much like single cells in an analogous gradient-free microenvironment (GFA). More interestingly, we found that cells are accelerated along the outward-pointing gradient of a spheroid and that this results in faster and more directed migration patterns.

Our results are consistent with persistent motion, as U87-MG cells tend to have aligned velocities at short time scales which means they follow a preferred direction of motion with persistence; given by *P* > 0. However at long time lags, motion is uncorrelated i.e. persistence is lost and cells move more diffusely. This may arise from the collisions and changes in the cells’ environment such as chemicals and extracellular matrix that is being constantly remodeled. Therefore, the persistence observed for cells in the GFA is captured by the PRW model, which describes the motion of a self-propelled cell migrating along a preferred axis. However, this model fails to describe the gradient force exerted by the spheroid. So here we showed that the induced bias drives the cells’ motion along the radial axis and increases their superdiffusivity. Assuming that the spheroid only has an external influence on cell migration – i.e. its presence changes the external environment but has no intrinsic effect on the cell – the effect of the spheroid can be modeled by adding an external bias term in the Ornstein-Uhlenbeck process. The resulting model is a biased PRW (BPRW) model whose bias amplitude decreases with the distance from the spheroid^17^.

This radial bias might arise from chemical gradients (nutrients, chemo-attractants or repellents), cell density gradient, or from the reorganization of the matrix network as the cells move through it^13–15^. The nature of such cell-matrix reorganization includes irreversible proteolysis^18, 19^. We also find indications of this behavior, however, the loss of anisotropy over time suggests that the open paths could be closing again for large time delays. Therefore despite the cells’ persistence, the close interplay with the local environment will gradually stray the cells from their primary directions of motion.

Although the anisotropic PRW (APRW) model has proven to be an accurate description of 3D cancer cell migration in gradient-free environments^16^, we chose to use the simpler PRW model – which still captures the most important features of cell migration – as the basis for our BPRW model.

Even though model systems like matrix-embedded spheroids have greatly helped our understanding of cancer metastasis, they cannot recreate the entire metastatic process. Therefore, we limited our study to the initial invasion of the local microenvironment during which we identified a strong and space-dependent repulsive force on migrating cells caused by the cancer tumor. Further experiments should be conducted to estimate the cues and identify more specifically the nature and chemical origin(s) of the bias. One possible direction would be to mimic gradients of cell density or chemical factors as was previously done for breast cancer cells in a growth factor gradient^20^. This approach is also promising for large drug-screening processes^21, 22^ and development of new anti-cancer strategies.

## Materials and methods

### Cell culture

Uppsala 87 Malignant Glioma (U87-MG) cell line was cultured in Dulbecco’s Modified Eagle Medium (DMEM) with 10% Fetal Bovine Serum (FBS) and 1% penicillin/streptomycin (Gibco) at 37°C and 5% CO_2_ and harvested at 90% confluency.

### Single cell migration assay - GFA

In ultra-low attachment 96-well flat-bottom microtiter plates (Corning), ~ 3,500 cells pr. well were embedded in a final volume of 100 μL of 65% Matrigel™ (Fischer Scientific) mixture in medium. Prior to mixing, Matrigel™ was thawed on ice (or overnight at 4°C) and pipette tips were stored at −20°C. Furthermore, all experiments in this study were performed using the same batch of Matrigel™ to account for natural variations. After mixing cell culture and Matrigel™ mixture, the plate was incubated (37°C and 5% CO_2_) for ~1 hour until it had solidified completely. Then 100 μL medium was added to each well to ensure plentiful nutrient supplies and avoid condensation on the inside of the plate cover when imaging.

### Spheroid invasion assay - GA

Tumor spheroids were formed by suspending ~ 325 cells in 200 μL cell culture medium pr. well in a 96-well ultra-low attachment U-bottom plate (Corning) and incubated (37°C and 5% CO_2_) for ~ 72 hours. At this point, spheroids had reached a diameter of about ~80 μm. Then the spheroids were embedded in 65% Matrigel™ by aspirating 120 μL medium and adding 150μL ice cold Matrigel™ as detailed in^9, 23^. The plates were incubated (37°C and 5% CO_2_) to solidify completely (~1 hour) before adding 100 μL cell culture medium pr. well.

### Imaging

The Matrigel™ -embedded cells and spheroids in flat-bottomed or U-bottom 96-well plates were time-lapse-imaged using a JuLI™ stage real-time cell history recorder (NanoEntek, South Korea) placed inside an incubator (37°C and 5% CO_2_). Using the built-in software, exposure time, brightness, and focus were adjusted for each well. Time-lapse bright-field images were obtained using the fully automated x-y-z stage, every 3 min (GA) or 5 min (GFA) for at least 24 hours using a 10x objective to obtain a resolution of 440μm/pixel. The recorded images are 2D projections of a 3D migration. If a cell migrates too far away from the focal plane it is not visible anymore. The same goes if the cell migrates outside the field of view^16, 20^.

### Image analysis

To aid image analysis, we obtained binary masks of the raw images using the machine-learning-based detection algorithm Ilastik^24^. These masks were used as input for the TrackMate plugin^25^ for Fiji^26^ which extracted the trajectory, [*x*(*t*), *y*(*t*)], for each cell’s center of mass in a given time-lapse. We subsequently performed manual editing of the tracks with TrackScheme, as TrackMate might falsely detect or miss a few number of steps or links. Following the criteria discussed in ref.^27^, we terminated tracks when cells had migrated either i) along the edge of the image for more than 15 min, ii) less than 50 μm over their entire trajectory to remove the contribution from immobile/dead cells, as well as when cells either iii) divided or iv) merged. The quality and reliability of cell motility analysis is highly dependent on the image acquisition procedures. As we used a motorized stage during image acquisition, the vibrations or drift of that stage will translate into apparent cell movements. Therefore, we used plastic beads as phantom cells in ECM to evaluate the vibration noise. We concluded that the stage vibration is negligible, so the measured cell trajectories were directly used for the analysis without correction.

### Mean-squared displacement fitting

In order to characterize migration, we measured the diffusion via the time-averaged squared distance (msd) traveled by a cell over some time lag, *τ*:

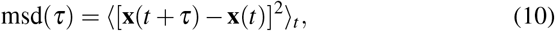

where **x**(*t*) is the time-dependent position vector and 〈…〉_*t*_ indicates an average over the entire cell trajectory.

We fitted the msd with eq. 2 to obtain the corresponding set of persistence time, *P*, and average speed, *S*, for each cell. Expectation values were obtained by fitting the resulting distributions (GFA, GA) with a log-Gaussian (fig. 2B) or Gaussian (fig. 2C), respectively.

The anomalous exponents, α, was found by fitting the ensemble averaged msd’s (MSD) with a power law as in eq. 1. Because these fits use non-linear least square methods, the results are highly dependent on the weighting function which compensates for the loss of resolution in the computation at large *τ*^28^:

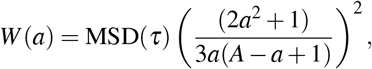

where *τ* is the time lag, as in eq. 10 and *a* = {1,2,3,…,*A*}is the number of points used for the fit, where we chose *A* to be 25% of the trajectory length to optimize the fits and discarded all fits with *R*^2^ < 0.99, as detailed in^9^ and references therein.

### Trajectory simulations

The BPRW simulation is described by a normalized Langevin equation, which provides the velocity change between two successive steps 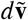. For the BPRW, the Langevin is the sum of the resistance to motion (first term), a time-dependent bias (second term), and random fluctuations (third term)^29^:

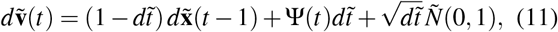

where 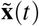 is the normalized position vector, 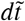 is the normalized time step, **Ψ**(*t*) the bias induced by the spheroid and 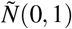 a scalar drawn from the standard normal distribution.

In order to get the normalized Langevin (eq. 11), the position vector is normalized as

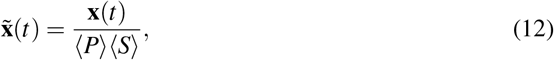

where the mean persistence time, 〈*P*〉, and migration speed, 〈*S*〉, are averaged over all experimentally-obtained cell trajectories for either the spheroid invasion assay (GA) or the single-cell migration assay (GFA); as detailed above.

The time step is re-scaled as:

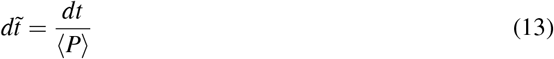

and the resulting trajectories have steps of

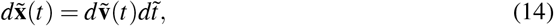

such that the position vector at time *t* becomes

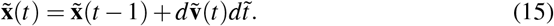

In the homogeneous environment experienced by migrating cells in the GFA, we assume that the bias term is null (**Ψ** = 0) and the model in eq. 11 reduces to a PRW model. In contrast in the GA case, we assume that the spheroid induces a bias (**Ψ** ≠ 0). This bias is presumably repulsive, radial and negligible far from the spheroid. Therefore, we hypothesize that the space-dependent field is proportional to −1/|**r**|, where **r** is the radial vector pointing outwards from the spheroid’s center to the cell position. We also assume the bias to be repulsive i.e the gradient is largest when *d***x**(*t* – 1) is antiparallel to **r**(*t*). The resulting time-dependent drift is therefore described by:

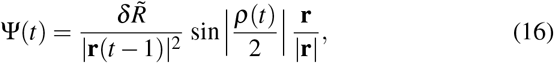

where *δ* is a dimensionless responsiveness to the bias, **r**/|**r**| is the unit radial vector, 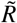 is the re-scaled radius of the spheroid, *R*_0_, such that

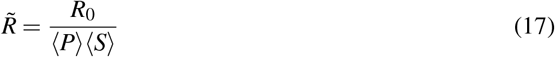

and *ρ* (*t*) is the angle between **r**(*t*) and the displacement vector, *d***x**(*t* – 1):

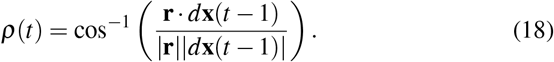

Both the direction and amplitude of the bias were inferred from the experimental results, as well as 〈*P*〉 and 〈*S*〉. Hence, *δ* was found by comparing the ensemble-averaged radial velocity in eq. 9 of data with the ensemble-average of *N* = 1,000 simulations of trajectories described by eq. 12–18, with *dt* = 0.5 min, *n* = 800 steps and a minimal displacement, *d* = 50*μ*m to remove the contribution from immobile/dead cells. Then, by minimizing the normalized root-mean-squared error we estimated the bias amplitude to be *δ* = 3. Taking all this together, trajectories for a BPRW and PRW were obtained with the equations above, with N=1,500, *n* = 800, *dt* = 0.5min and the following model parameters:

## Author contribution statement

MTS did the spheroid imaging (GA), MA the single-cell imaging (GFA) and LJ supervised the project. All authors conceived the idea, developed the mathematical models and discussed the results. Likewise, all authors contributed and approved the final manuscript.

## Additional Information

The authors declare no competing interests.

