## Supplemental Figures for "Tumor spheroids accelerate persistently invading cancer cells"

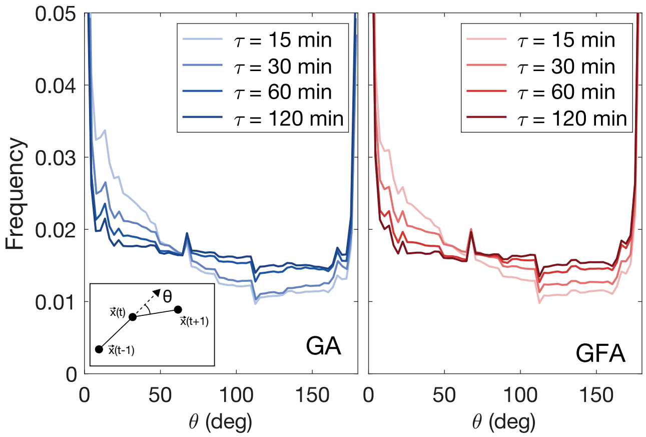


**Fig. S1** Normalized and smoothed distribution of turning angles, $\theta$, for GA (N=177) and GFA (N=305). The insert is a sketch of $\theta$ as defined in the main article. In both the GA and GFA cases, the distribution has local maximum at $\theta = 0$ and $\theta = 180°$and flattens for larger delays, $\tau$.

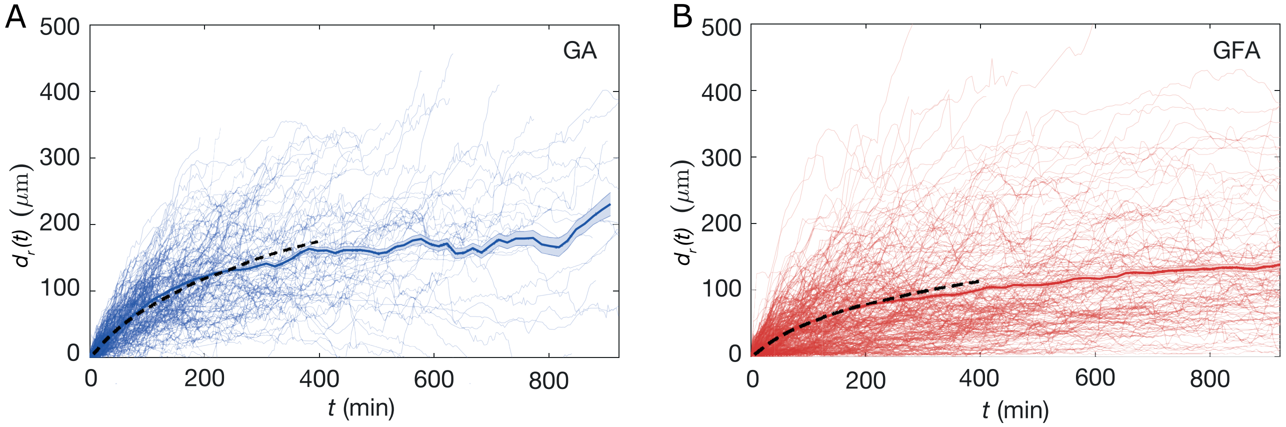


**Fig. S2** Time evolution of the average migration displacement, $d(t)$, versus time. **(A)** The thin lines represents individual trajectories (N=177) and the full line is the average distance and shaded area is one SEM. The dotted black line corresponds to the average simulated trajectories using the BPRW model. **(B)** The thin lines represents all tracked trajectories (N=305) and the full line is the average distance and shaded area is one SEM. The dotted black line corresponds to the average simulated trajectories using the PRW model.

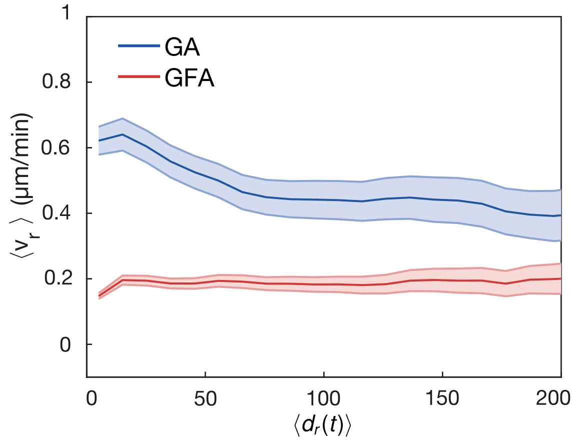


**Fig. S3** Evolution of the ensemble-averaged radial velocity, $\langle v_r \rangle$, (mean ± SEM) against radial distance, $d_r(t)$ for both GA (blue) and GFA (red). $v_r$ is binned on 15 µm with a moving average of 10 points.
